# Tools for Visualizing and Analyzing Fourier Space Sampling in Cryo-EM

**DOI:** 10.1101/2020.06.08.140863

**Authors:** Philip R. Baldwin, Dmitry Lyumkis

## Abstract

A complete understanding of how an orientation distribution contributes to a cryo-EM reconstruction remains lacking. It is necessary to begin critically assessing the set of views to gain an understanding of its effect on experimental reconstructions. Toward that end, we recently suggested that the type of orientation distribution may alter resolution measures in a systematic manner. We introduced the sampling compensation factor (SCF), which incorporates how the collection geometry might change the spectral signal-to-noise ratio (SSNR), irrespective of the other experimental aspects. We show here that knowledge of the sampling restricted to spherical surfaces of sufficiently large radii in Fourier space is equivalent to knowledge of the set of projection views. Moreover, the SCF geometrical factor may be calculated from one such surface. To aid cryo-EM researchers, we developed a graphical user interface (GUI) tool that evaluates experimental orientation distributions. The GUI returns plots of projection directions, sampling constrained to the surface of a sphere, the SCF value, the fraction of the empty region of Fourier space, and a histogram of the sampling values over the points on a sphere. Finally, a fixed tilt angle may be incorporated to determine how tilting the grid during collection may improve the distribution of views and Fourier space sampling. We advocate this simple conception of sampling and the use of such tools as a complement to the distribution of views to capture the different aspects of the effect of projection directions on cryo-EM reconstructions.

## 1. Introduction

Cryo-EM reconstructions using single-particle methods are appearing at an ever-increasing rate, and the number of map depositions into the electron microscopy database is growing accordingly [1-3]. Advances to technology are continuing to propel the field and open up directions for numerous biological applications [4]. However, there remains a persistent problem. Current vitrification technologies for preparing samples for imaging lead to adherence of macromolecular specimens to one of the two air-water interfaces [5, 6]. This seems to affect the vast majority of specimens that are currently analyzed using cryo-EM [6]. The interfaces attract hydrophobic patches on macromolecular surfaces, and macromolecules therefore adopt “preferred orientations” with respect to the electron beam. One, or several, preferred orientations can exist within a given dataset. This results in a non-uniform projection orientation distribution within a cryo-EM dataset.

The orientation of a macromolecular object embedded within a vitreous ice layer and imaged using an electron microscope can be described by three Euler angles, ϕ, θ, and Ψ. In conventional notation, the three Euler angles describe: ϕ, the rotation of the object around the Z axis, or the longitudinal position around the zenith of a sphere; θ, the rotation of the object around the *new* Y axis, or the latitudinal position about a sphere; Ψ, the rotation of the object around the *new* Z-axis, or the relative in-plane rotation [7]. For each image, the angles ϕ and θ uniquely describe the relative position of the object on a sphere. A scatter plot of ϕ versus θ angles provides a complete description of the set of orientations assigned to the object within a cryo-EM dataset. Therefore, the standard manner by which to represent an orientation distribution of a set of images in cryo-EM is to plot Euler angles ϕ and θ, and to assess their relative distribution.

We and others described efforts to provide a quantitative measure to a given orientation distribution [8, 9]. An uneven set of projection views compromises the cryo-EM reconstruction. During the reconstruction procedure, each projection is inserted as a central slice through the 3D Fourier transform of the object orthogonal to the direction of the projection [10]. Therefore, non-uniformity in the set of projection distributions also leads to non-uniformity in Fourier space sampling [8]. We refer to “sampling” simply as the number of times that a representative point in Fourier space has been measured. There are two typical cases to consider during the reconstruction process: (1) the case where all Fourier voxels receive adequately high sampling and (2) the case where certain sets of contiguous voxels have not been sampled at all.

The first case occurs, for example, when projections are distributed azimuthally around the imaged object. For a C/D-fold symmetric object, or for helical assemblies, a regular distribution of projections about the object and perpendicular to the symmetry axis is commonly referred to as “side-like” views. Modulations in the regularity of these views (i.e. modulating the ϕ angle) produces a set of side-like distributions, where there are less well-sampled pockets of Fourier space. A typical second case occurs when projections do not (approximately) form any great circle around the imaged object and there are no additional orientations. In cryo-EM, the second scenario is commonly observed for asymmetric objects that have a single preferential orientation; it can also be observed for symmetric objects, but depends on the symmetry and the location of preferred orientation with respect to the symmetry axis. This scenario leads to unsampled zero values in the transform. Numerous synthetic case studies of both fully sampled distributions and those containing missing information have been recently described [8]. For all the above cases, the set of projection orientations are by definition non-uniform, leading to uneven angular sampling of Fourier voxels. The non-uniformity of orientation distributions, and the concomitant irregularity of the sampling, was shown to attenuate global resolution. For cases characterized by zeros, there will also be a limit to the maximum attainable resolution, even as the number of particles increases.

Attenuation of global resolution due to non-uniformity can be accounted for by a geometrical factor that we previously termed the sampling compensation factor (SCF) [8]. The name derives from the idea that the non-uniformity in sampling leads (approximately) to a non-uniformity in effective noise variance, but otherwise does not affect the envelopes describing the reconstruction. The SCF is related to the average of the reciprocal sampling over shells in Fourier space. The reciprocal sampling arises naturally in calculations where noise is regrouped and effectively decremented in performing direct Fourier reconstructions. An underlying idea is that Fourier space filling more directly affects the signal-to-noise ratio; however, it is harder to interpret the effect on the SSNR directly from the set of projection views. Also, we pointed out that when there is missing data, it is necessary to assign appropriate variance to unmeasured values of the transform. This leads to a more appropriately defined version of the SSNR, which accounts for zeros, and is termed SCF*. This SCF* thus accounts for the decrement of the corrected SSNR due to the collection. Under control experiments, and assuming that orientations are perfectly established, the SCF and SCF* appear to be adequate estimates for how resolution is attenuated due to uneven sampling geometry.

Multiple relevant questions remain: how does a given orientation distribution translate into Fourier space coverage? Which regions are over-sampled? Which are under-sampled? If present, where are there zeros in the transform? In general, the relationship between a qualitative orientation distribution and a number, such as the SCF/SCF*, used to evaluate its “goodness” is not intuitive to the cryo-EM practitioner. Indeed, most users are hard-pressed to explain whether the set of views they have acquired are adequate for recapitulating structural details of the imaged object at a defined resolution.

To address current limitations, we developed a set of diagnostic tools to visualize Fourier space sampling. In this work we introduce a plot, which describes the sampling on the surface of a sphere as a representation of Fourier space coverage. We also developed a graphical user interface (GUI), which helps users visualize both the orientation distribution and Fourier space coverage side-by-side. The GUI facilitates understanding, in a more intuitive manner, how missing data and/or sampling modulations can affect the reconstruction. The GUI reports both SCF and SCF* values, which quantitatively describe the sampling distributions. The SCF* values, as opposed to SCF values, are adjusted for the presence of zeros in the transform and provide a more intuitive understanding of the effects of sampling on resolution attenuation. We present multiple case studies to illustrate the ideas. These tools should be generally relevant to the cryo-EM community and lead to a better and more quantitative understanding of the effects of preferred specimen orientation and non-uniform projection distributions.

## 2. Results

### 2.1. A description of the sampling plot constrained to the surface of a sphere

We wished to provide a framework that will allow the user to directly visualize the effects of a projection distribution on Fourier space sampling. To this end, we developed surface sampling plots, which provide a quantitative description of Fourier space coverage. Our conception of sampling is as follows. Every point of Fourier space may or may not be measured by a given projection. For each lattice site included in the sampling map, we measure the total number of times that the region indicated by the point is measured by the set of projections. We can use a set of lattice sites that are constrained to a surface of a given Fourier radius, which is our sampling map constrained to a (hemi)sphere. Since the direction of a projection is orthogonal to the 2D slice within the 3D transform of the object, an increase in the number of projections along a particular view accordingly populates the surface sampling plots orthogonal to the direction of the projection. The specific quantitative considerations for the projection plots and the surface sampling plots are discussed in sections 3.1 – 3.6. The considerations when applying symmetry or an arbitrary tilt angle are described in sections 3.7 – 3.8. The effects of different orientation distributions on surface sampling plots will be evident in the subsequent section.

### 2.2. Examples of sampling plots for different orientation distributions

To determine how different projection orientation distributions contribute to surface sampling plots, we examined multiple cases that arise in experimental settings. Five different orientation distributions are demonstrated: (1) uniform, (2) top-complement, (3) side-like, (4) side-modulated, and (5) top-like. We previously examined these distributions when we introduced the SCF [8]. The first four cases represent fully sampled distributions, wherein every Fourier voxel has been assigned some value during the reconstruction procedure. The last case, the “top-like” distribution, results in a missing cone of information, the presence of zeros in the transform, limits the attainable global resolution (as reported by the SSNR/FSC), and as described previously, must be treated distinctly [8]. These distributions will now be examined using surface sampling plots.

#### 2.2.1. Uniform distribution

The uniform distribution describes a scenario wherein all angular orientations corresponding to the imaged object are present within the dataset with (approximately) equal occupancy. In an experimental setting, it occurs when there is a complete absence of preferential orientation, and the specimen is randomly distributed within the ice layer. All Fourier voxels on a single shell are (approximately) equally sampled. This even sampling is evident within the surface sampling plots in Figure 1A. The uniform distribution is characterized by an SCF/SCF* value of 1. It is an ideal case devoid of any anisotropy and characterized by the maximum attainable global resolution for the given experimental envelope.

**Figure 1.**
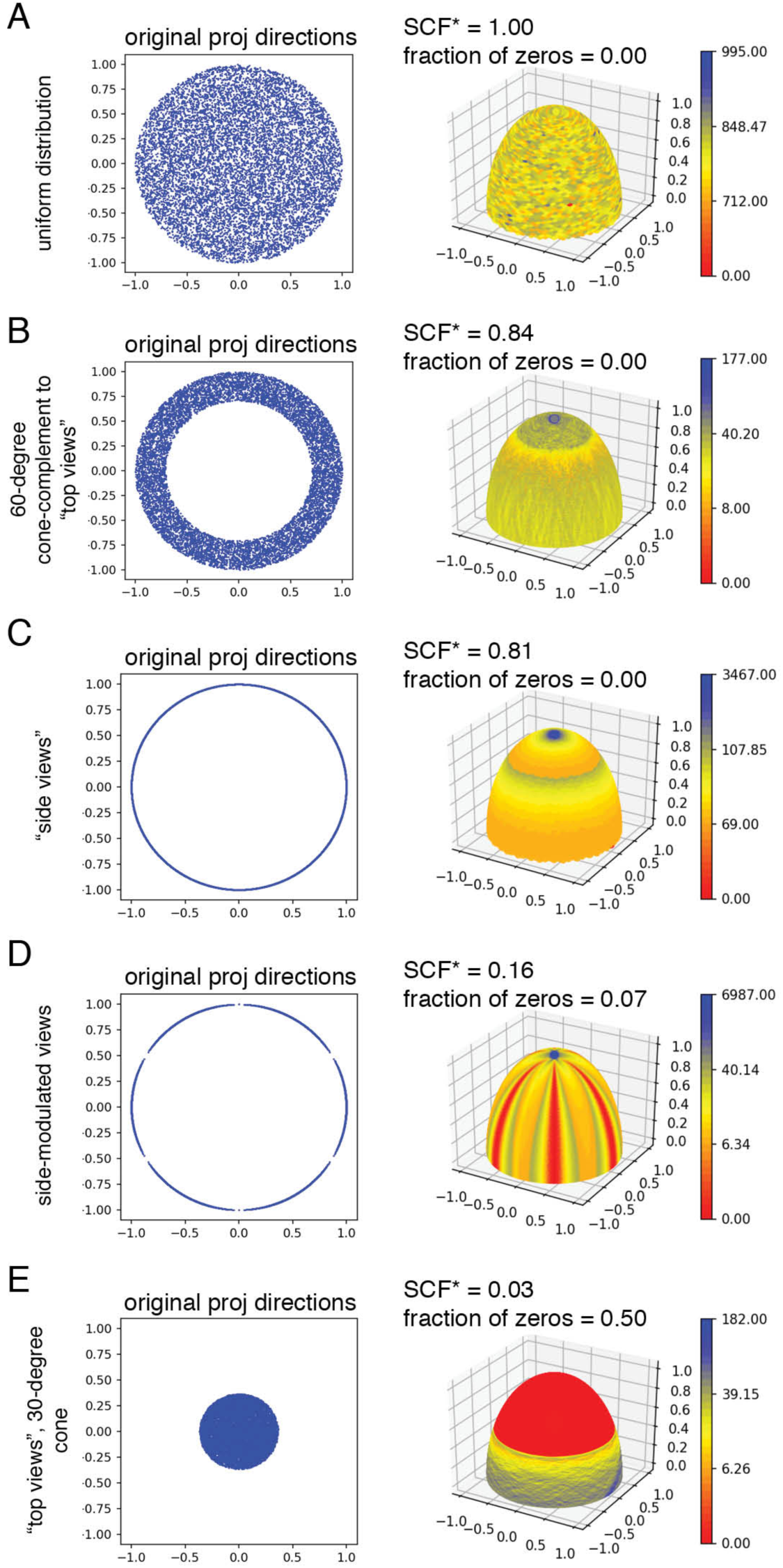
Evaluation of Fourier space sampling for distinct orientation distributions. Five different datasets taken from [8] are analyzed, each of which is generated from 10,000 projection images. The panels show (left) the projection distribution and (right) the surface sampling plot for a particular dataset. Values along the axes indicate directions on the unit sphere. On the surface sampling plots, the scale bar refers to a histogram of sampling values. Multiple different types of distributions are shown, ordered by the extent of sampling. The datasets correspond to: (A) uniform distribution, (B) Complement to “top-like” views, (C) pure “side-like” views, (D) “side-modulated” views, and (E) “top-like” views.

#### 2.2.2. Top-complement distribution

The top-complement distribution describes a scenario wherein all angular orientations of the imaged object are present within the dataset with (approximately) equal occupancy, with the exception of projections with θ-angle (tilt) constrained, so that no projection is taken interior to the cone with half angle X. More simply, all orientations are sampled, but there are no “top-like” views (top of projection sphere, equivalently north pole). In an experimental setting, the top-complement distribution may arise for rotationally symmetric objects that are characterized by a preferential orientation along their side, but not along their top views, and with a distribution of intermediate tilt angles about the side. The proteasome is one such example. For such distributions, there is an increased sampling along the Z-direction compared to the X/Y plane, and the increase is directly proportional to the size of the cone. This can be observed within the surface plots. For example, for an object characterized by a 60°-cone-complement (i.e. the θ-angle is randomly modulated to 60°<θ<120° or +/-30° above the equator), there is an approximately two-fold increase in sampling toward the Z-axis in comparison to the X/Y axes (Figure 1B). The top-complement distribution is characterized by SCF/SCF* values that range from 8/π^2^ (∼0.81) toward a maximum of 1, where 8/π^2^ is the theoretical value that has been described previously for perfect side views [8]. Top-complement distributions are some of the best-sampled cases that can occur within an experimental setting.

#### 2.2.3. Side-like distribution

The side-like distribution describes a scenario wherein the θ angle is restricted to a value of 90°, or along the equator, but with an otherwise uniform ϕ distribution. For the case of side views, it is well known that the sampling is complete [11]. In an experimental setting, the side-like distribution is most closely associated with helical specimens [10]. For helical specimens (especially long filaments), there is often a very small or negligible deviation of the θ angle from the equator. The side-distribution may also be associated with rotationally symmetric single particles that adopt a preferred orientation to their side, but not their top views. However, such cases typically have some amount of tilt angle modulation about the equator, and therefore may be better characterized by a top-complement distribution with a large cone. Again, there is a large accrual of sampling along the Z-axis compared to the X/Y plane. However, the extent is more pronounced than for the top-complement case. For a perfect side distribution, the accrual of sampling can range by up to an order of magnitude (Figure 1C). The SCF for the perfect side distribution evaluates to the theoretical value of 8/*π*^2^ = .81 reported in [8] and the SCF* is identical to SCF, because there are no zeros. In the side-like cases, the residual non-uniformity compromises the SSNR by ∼20% in comparison to uniform, and would therefore have a small effect on experimental resolution attenuation.

#### 2.2.4. Side-modulated distribution

The side-modulated distribution is an extension of the side-like distribution and describes a scenario wherein the θ-angle is restricted to a value of 90°, but the ϕ-angle is modulated to varying extents. In an experimental setting, the side-modulated distribution can arise for objects characterized by low rotational symmetry (e.g. C3, C4, etc.) and adhered to the air-water interface along their side view (sampling for a C2-symmetric object, adhered to its side view, behaves like a top-like distribution described below, as for example the HIV intasomes) [12]. The higher the rotational symmetry, the closer the behavior will be to perfect side views. Similar to the side-like distribution, there is an increased sampling along the Z-direction at the expense of the X/Y plane. In addition, there are also pockets of incomplete sampling around the perimeter. Depending on the modulation of the ϕ-angle, these pockets can be substantial. For example, when there are three full periods of ϕ-angle modulation (corresponding to C3 symmetry), the values within the gaps may closely approach zero (Figure 1D). In the most extreme cases, especially when the dataset is small (<10,000 particles), there could be zeros within the gaps in the transform. The side-modulated cases are characterized by SCF values ranging from 0 < X < 8/π^2^. X depends on the symmetry of the object, and the extent of ϕ-angle modulation. For low-symmetry objects with small ϕ-angle modulation, the values can be in the range of ∼0.1-0.2. The situation is quickly ameliorated with higher rotational symmetry or lower modulation, as previously demonstrated [8]. Assuming that the dataset size is sufficiently large, the side-modulated case would almost always be fully sampled. Nonetheless, the resolution may be attenuated by up to an order of magnitude, depending on the amount of modulation.

#### 2.2.5. Top-like distribution

The top-like distribution is a pathological distribution that describes a scenario characterized by a single dominant view. It is one of the most common distributions encountered within experimental cryo-EM datasets, due to the problem of preferred orientation [6]. It arises because the specimen adheres to one of two air-water interfaces (AWI) in a single orientation (with a random in-plane distribution), giving rise to a conical distribution of projection views. The top-like distribution is characteristic of any asymmetric object that adheres to the AWI in a single orientation. It can also describe the sampling for rotationally symmetric objects that adhere to the AWI along their top, but not their side interfaces, or for C2-symmetric objects adhered to their side view. This is a pathological distribution that is, in the absence of other views, characterized by zeros in the transform. For a 30° cone, one can readily observe several features to such distributions (Figure 1E). First, there is increased sampling closer to the X/Y plane. The magnitude can vary significantly, depending on the size of the cone. Second, there are zeros in the transform, indicated by red regions on the plots. As described previously, the zeros need to be treated differently; unit variances must be assigned to unmeasured voxels when calculating the SSNR [8]. The value that gives the decrement of the appropriately defined version of SSNR, due to the type of sampling, is therefore reported as SCF*. Indeed, distributions that have primarily one view, can have SCF* values that differ by up to three orders of magnitude [8]. There is also a finite resolution limit that depends on the size of the cone and the proportion of sampled/unsampled voxels. Top-like distributions can easily be improved through the addition of alternative views. Given their pathology, even small modifications to the orientation distribution can have a large effect on Fourier space sampling and global resolution. The sampling plots, the resulting SCF* values, and the numerical assessment of zeros provide an important insight to the extent of this pathology.

### 2.3. The effects of tilting the grid on the sampling distribution and the SCF value

One of the motivations for developing sampling plots as a readout of Fourier space coverage was to provide a quantitative framework with which to evaluate the effects of altering an experimental orientation distribution. For example, it is known that an orientation distribution can be altered through physical tilting of the microscope stage during the imaging experiment [13, 14]. To this end, we examined how different orientation distributions, and the corresponding SCF values, would be affected by applying a nominal tilt angle. For each distribution described in figure 2, we applied a transform to simulate a 30°-tilt or a 45°-tilt angle inside the electron microscope (see section 3.8 for further details).

**Figure 2.**
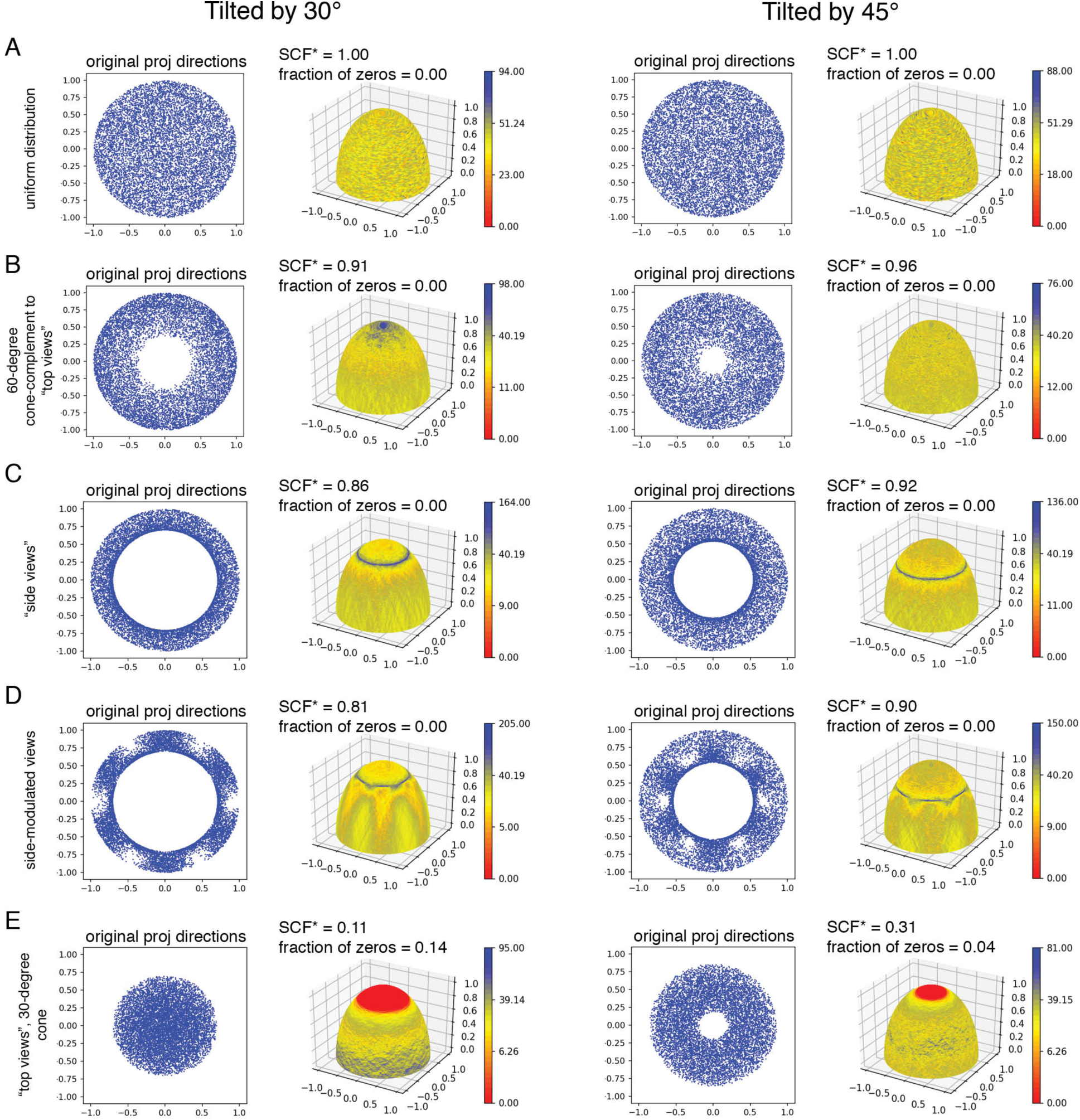
Evaluation of Fourier space sampling for distinct orientation distributions, tilted by 30° and 45°. A tilt angle was applied to each of the five datasets shown in Figure 1. The panels show (left) the projection distribution and (right) the surface sampling plot after applying either a 30° or a 45° tilt angle to each dataset.

The uniform distribution is not affected by the application of a tilt angle, as would be expected. Accordingly, no changes are observed to the sampling distribution or the SCF* value (Figure 2A). For the top-complement case, the application of a tilt angle spreads out the distribution to a nearly uniform Fourier coverage. The SCF* increases from 0.84 to 0.91 for the 30° tilt and to 0.96 for the 45° tilt. The increased uniformity is evident both in the new sampling plot and the new projection distribution (Figure 2B). For the perfect side case, a tilt angle converts the distribution into one that behaves similar to the top-complement scenario. The projections are pushed from a single thin band around the equator to a thicker band, due to the tilt. The peak of the sampling down the z-axis is pushed to a ring encircling the z-axis, which directly relates to the tilt angle. A 30° tilt angle improves the SCF from 0.81 to 0.86, and a 45° tilt angle improves to 0.92 (Figure 2C). A similar situation is encountered in the side-modulated case. However, now the SCF improves more substantially, from 0.07 to 0.81 and 0.90 for the 30° and 45° tilts, respectively (Figure 2D). This implies that, with certain low-symmetry distributions, the application of a small tilt angle may have substantial effects to improve the quality of the map. The largest improvements to sampling arise for the cases with missing data. Notably, when there is missing data, the SCF can be very small, ranging by up to three orders of magnitude [8]. The introduction of additional orientations can substantially improve the sampling. For example, in the case of a 60-degree missing cone, the application of a 30° tilt angle improves the SCF* by over five-fold, while a 45° tilt angle improves by fifteen-fold (Figure 2E). This means that simply tilting the grid can have substantial benefits to improving the reconstruction. Indeed, many experimental cases have already made practical use of tilts to solve the preferred orientation problem and resolve the structure of a macromolecular object of interest [14-21], including samples as small as 40 kDa [22], and as large as the ∼100 mDa nuclear pore complex [23, 24]. The sampling plots, and the SCF* values, now provide quantitative insight explaining the benefits observed through experiment.

### 2.4. The effects of symmetry on the sampling distribution and the SCF value

Symmetry has the effect of increasing the number of views. For single-particle datasets, symmetry multiplies the effective dataset size, and can also improve the SCF* by increasing Fourier space coverage. It is worthwhile to distinguish these effects. The first effect is straightforward: an *n*-fold symmetry effectively multiplies the dataset size by *n*. The second needs to be considered more closely. Whereas preferred orientation affects both symmetric and asymmetric samples, symmetry can mitigate the sampling problem. An example is the 20S proteasome, (EMPIAR-10025), displayed in Figure 3. This specimen has two preferred orientations within a region of the asymmetric unit – one along its “side” view and one along its “top” view. Due to D7 symmetry, both orientation are equally likely to be observed in one of fourteen different positions in the usual representation of projections on the sphere. Consequently, these two preferred orientations become repeated multiple times when the object is reconstructed, and the resulting SCF* value improves from 0.04 to 0.84, by a factor of ∼20 (Figure 3A-B). However, just because an object has symmetry does not mean that symmetrization will lead to a more isotropic reconstruction. The isotropy depends on the axis of preferred orientation and its relationship to the molecular symmetry. This is demonstrated when the symmetry for the proteasome is split between its constituent side and top views. If the axis of preferred orientation is positioned perpendicular to the seven-fold molecular symmetry axis (i.e. along the “side-like” views), then symmetrization improves the SCF* by virtue of improving Fourier coverage – along X/Y and Z axes (Figure 3C-D). In contrast, when the top views are symmetrized, Fourier slices are inserted proximal to the X/Y plane. Applying symmetry to the reconstruction merely leads to hyper-sampling near the X/Y plane of the transform, but not along the Z-direction. In this case, symmetry does not benefit the reconstruction at all, and there is virtually no gain in the SCF (Figure 3E-F). Therefore, most of the gains in the SCF* value of the proteasome arises from the contributions of the side-like views.

**Figure 3.**
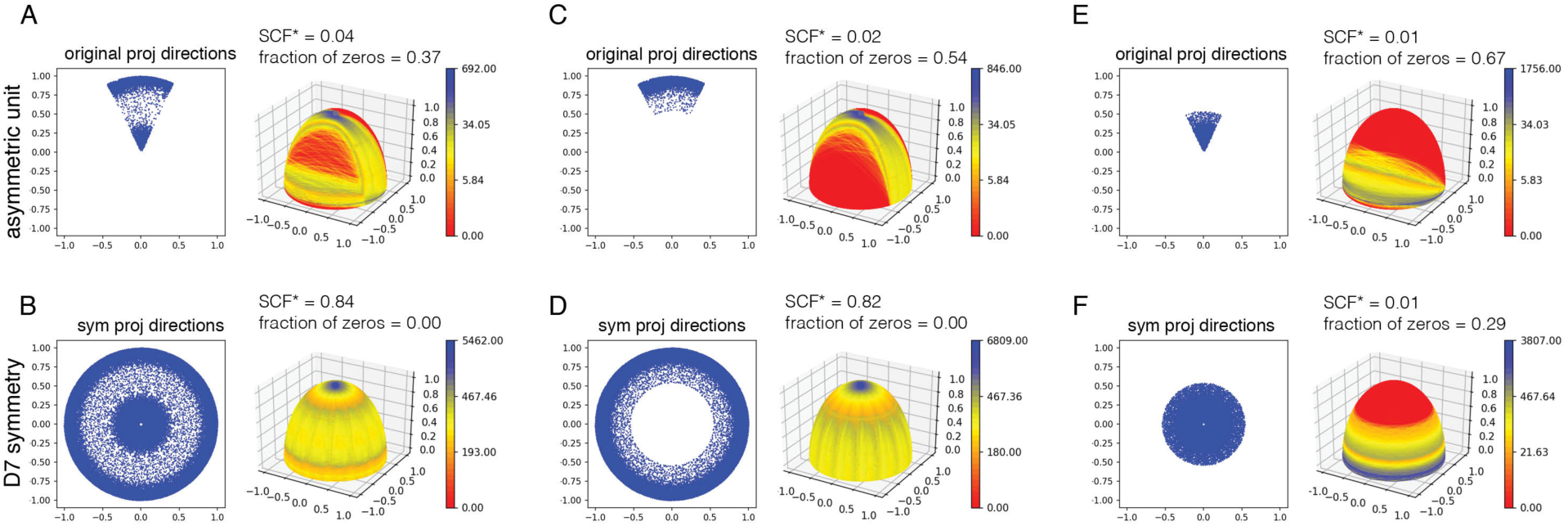
The effect of symmetry on sampling distributions – proteasome dataset from EMPIAR 10025. The panels show (left) the projection distribution and (right) the surface sampling plot for each dataset. (A) Euler angles corresponding to a single asymmetric unit for the proteasome dataset. (B) Symmetrization of the distribution in A using D7 symmetry. (C) Euler angles corresponding to a single asymmetric unit of the proteasome, where the θ angle is restricted to the range 45° < θ < 90°, generating “side-like” (or top-complement) views. (D) Symmetrization of the distribution in C using D7 symmetry. (E) Euler angles corresponding to a single asymmetric unit of the proteasome, where the θ angle is restricted to the range 0° < θ < 45°, generating “top-like” views. (F) Symmetrization of the distribution in E using D7 symmetry.

The presence of symmetry also simplifies the angular search space under experimental conditions. Specifically, consider two equivalent scenarios: (1) assigning angles to a single asymmetric unit, then symmetrizing the reconstruction; (2) assigning angles to the entirety of the (hemi)sphere, then performing an asymmetric reconstruction. The resulting maps would be identical. However, the first scenario is computationally less expensive. Therefore, many algorithms will only search orientations within a confined space characterized by the point group symmetry. For such cases, it is necessary to symmetrize the orientations in order to properly determine the sampling and the related SCF values. Just like in the case with the 20S proteasome (Figure 3), particle orientations for apoferritin (EMPIAR-10216) are only assigned to one asymmetric unit, out of its eight symmetry-related copies. If symmetry is not specified, the resulting SCF* value is 0.03; however, when the proper octahedral symmetry is taken into account, the SCF is correctly determined at near unity (0.99), which is consistent with the isotropic reconstruction (Figure 4). Therefore, it is necessary for users to properly specify the appropriate symmetry to completely sample Fourier space when analyzing orientation distributions for their datasets.

**Figure 4.**
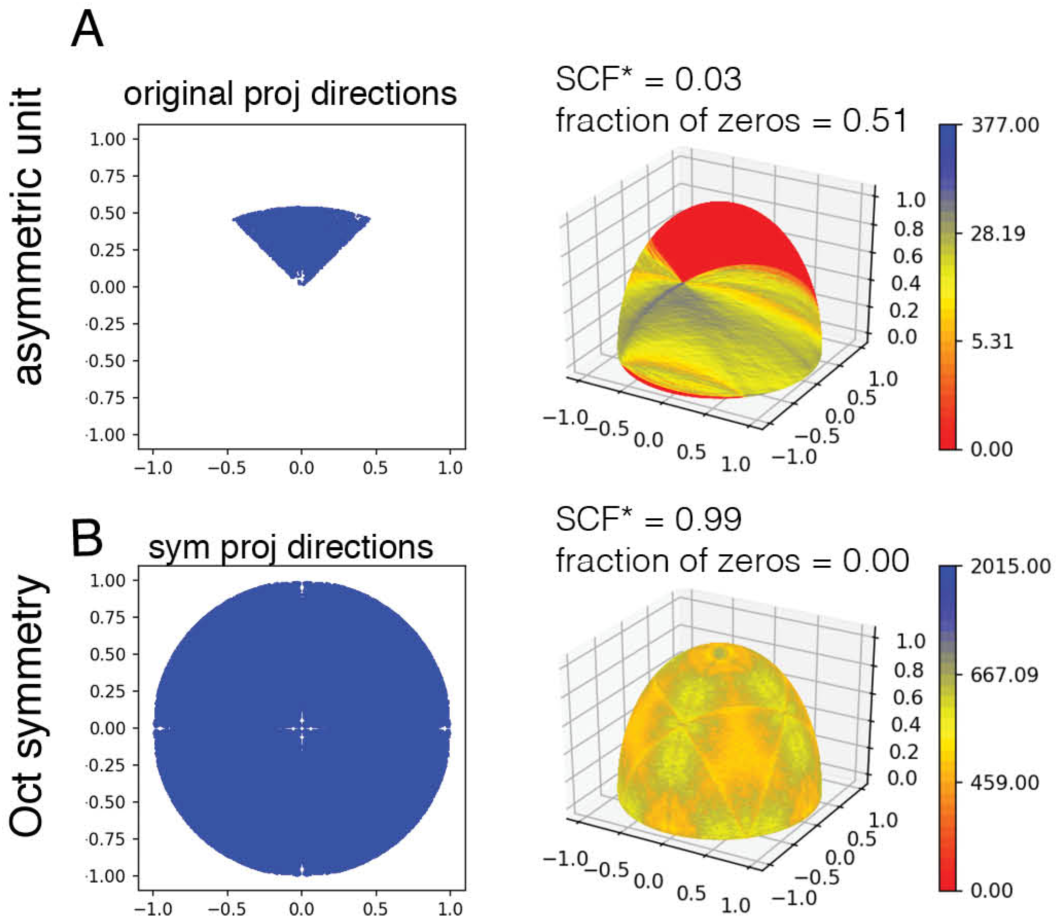
The effect of symmetry on sampling distributions – apoferritin dataset from EMPIAR 10216. The panels show (left) the projection distribution and (right) the surface sampling plot for the deposited dataset. (A) Euler angles corresponding to a single asymmetric unit for the apoferritin dataset. (B) Symmetrization of the distribution in A using octahedral symmetry.

As a general rule of thumb, SCF values greater than or equal to ∼0.81 (which corresponds to the SCF value of uniform side views and a fully sampled reconstruction) describe a situation with adequate sampling to faithfully recapitulate all the structural details of cryo-EM reconstructions. This particular value is taken from prior derivations, which show that ∼0.81, or 8/π^2^ represents the fully sampled case and is characteristic of perfect side views [8]. We thus suggest that this value can be considered the minimal goal for an experiment. For SCF values <0.81, one should consider whether adding views – e.g. by tilting – would contribute more to improving the resolution than any associated tradeoffs, such as thicker ice. More specific and generalizable recommendations are currently the subject of ongoing work.

### 2.5. Description of the GUI

The tools have been incorporated into a graphical user interface to make it easier for experimentalists to examine their orientation distributions. Figure 5 shows the current version of the GUI. The only required input from the user is a file containing the Euler angles Ψ, θ, and ϕ in 3 column format. This format can easily be converted from Frealign, [25] Cryosparc, [26] Relion, [27] or other programs using any convenient spreadsheet editor. The Ψ angle is required to determine the proper orientation upon application of a tilt angle (see sections 2.3 and 3.8). The program returns a surface sampling plot, an Euler distribution plot, a histogram profile of sampling values at the defined Fourier radius, and SCF and SCF* values. For cases with complete Fourier sampling, the SCF and SCF* values would be identical. For cases with unsampled regions of Fourier space, the SCF* value reports the properly corrected geometrical factor. The user is also given the option to change certain parameters, including the number of particles to use, the symmetry, and the Fourier radius. A default Fourier radius is provided to determine the SCF, which makes it possible to obtain a result within a few seconds on a standard processor. A larger radius (for example, ½ the boxsize) can be used, but should result in a similar sampling plot and pattern in the histogram of sampling values, although the fraction of zeros will differ (see section 3.4). Finally, the tilt angle may be selected by a slider. This option allows the user to determine how the orientation distribution will change when the data is collected at a tilt angle. Determining an optimal tilt angle is relevant for datasets that are characterized by non-uniform sampling. As demonstrated in Figure 2D-E, a small tilt angle can dramatically improve the SCF. It is therefore always worthwhile to consider how to improve the sampling distribution (e.g. by tilting the data) rather than to collect more particles. Further work will be required to fully understand all implications, including selecting the optimal tilt angles for experimental data characterized by different types of sampling distributions, assessing the experimental attenuation of collecting data at a tilt angle, and determining the contributions of false positive orientation assignments.

**Figure 5.**
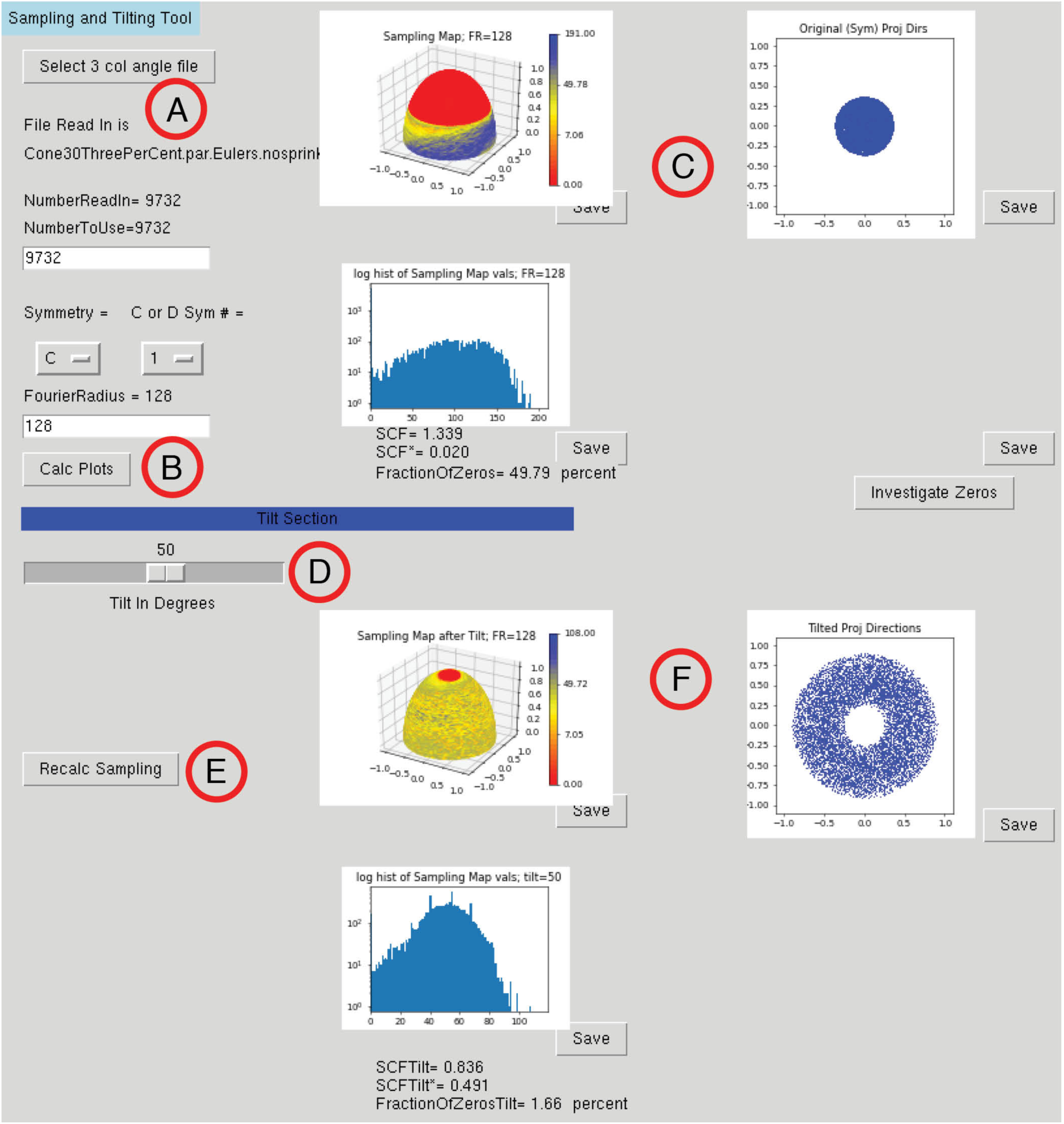
Screenshot of the graphical user interface (GUI) to evaluate sampling and the effects of tilting. Panels refer to the different options in the GUI and areas where the user has input: (A) the user must input a 3-column angle file and specify the number of particles, the symmetry, and the Fourier radius. (B) The “Calc Plots” button generates output plots. (C) Outputs include the surface sampling plot (top left), the original projection distribution (top right), and a histogram of sampling values for the Fourier voxels (bottom). The latter can be helpful to show, for example, the spread of sampling or how often values near zero occur. (D) A slider bar is used to assign a nominal tilt angle to the original projections. (E) The “Recalc Plots” button generates output plots after the application of a nominal tilt angle to the distribution. (F) The outputs here mimic those in C, except that each has been transformed in accordance to the specified tilt angle.

## 3. Mathematics related to sampling and the GUI

### 3.1. Plotting Projections in 2D

This section pertains to plotting of projections on a 2D plane. The purpose is to display the projections in a manner such that they can be viewed as a flat plot, and without projections bunching up near the equator (side views). In cryo-EM software, projections are conventionally plotted with the polar angle of the projection being the radial vector, and the azimuthal angle of the plot being identical to the azimuthal angle of the point on the projection sphere. This addresses an issue, that if simply *x, y* values of the projection were plotted, the points would appear to bunch up near the equator. To settle this issue in a principled way, one can seek a radial variable such that the measure on the resulting plotting plane transforms to the uniform measure on the sphere. That is, we seek *ρ*(*θ*) such that d*x* d*y* = (constant) sin *θ* d*θ* d*ϕ* (which is the uniform measure on the sphere), where (*x, y*) = *ρ* (*θ*) (cos *ϕ*, sin *ϕ*). As shown in the appendix, this leads to the unique solution 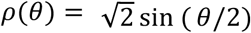, such that the maximum new “radius” = 1. This is how we make our 2d plots of the projections on the plane: the *x, y* values of the projection unit vectors are taken, the azimuthal angle is left intact, but the radial variable is changed, essentially from sin *θ* to 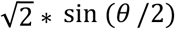. Over the interval [0, *π*/2] this function is hardly discernible from the simpler, 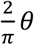; they agree at endpoints, and differ everywhere by less than 5%.

### 3.2. Projection-sampling transform

This sub-section discusses the linear relationship between the set of projection views and the sampling from a formal perspective. By casting this relationship into the appropriate basis, we argue that the set of projection views and the sampling restricted to a set of points on a sphere, contain equivalent information, by showing that the relationship is invertible (up to the accuracy determined by the spacing on this sphere). This is one of the main concepts underlying the current work. We achieve this by showing that the linear mapping has spectrum bounded in magnitude away from zero.

If 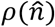 is the (possibly discrete) density of projections at the point on the unit sphere, then we define the sampling as done previously [8], 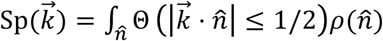, where Sp is the sampling and Θ is the indicator function, taking value of unity if the condition holds, and zero otherwise. This is the slab condition: at each lattice site 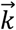, one counts the number of times the site belongs to the slab of points of unit width that is normal to each of the projection directions 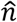: see Baldwin & Lyumkis [8] section 3. This is the most simplistic conception of the filling of Fourier space. The last relationship may be recast as an angular convolution over the surface of a fixed sphere:

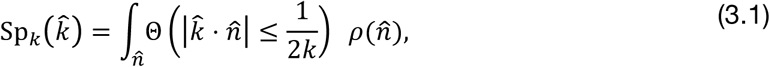

meaning that it is an equation of the form Output 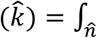 Linear Kernel 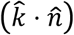 · Input(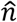). In the next sub-section, we show that this same linear kernel is invertible, in our numerical implementation. The sampling thins approximately as 1/2k. To see this, consider a 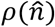 which describes a single projection. For a single projection and a unit sphere, 2*πk* will sample the circumference of the sphere out of 4*πk*^2^ available points on the surface area. The sampled points must be divided by the 4*πk*^2^ available points. The ratio of these quantities is 1/2k.

### 3.3. Invertibility of the transform

The equations of the form (3.1) can be brought to diagonal form, by means of a transformation to spherical harmonics (essentially the so-called Funk-Hecke theorem [28]). The resulting equation in the harmonic basis is:

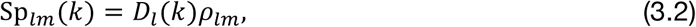

where *l* is the angular frequency and must be even, and *m* is the standard orientational parameter expressing the coefficients with respect to a special axis (commonly, *z*). Therefore *Sp*_*lm*_ (*k*) is the sampling and *ρ*_*lm*_ the projection density defined on the sphere in the harmonic basis, whereas *D*_*l*_ is the kernel of (3.1) in this basis (and does not depend on the label, *m*, only on the frequency, *l*). Approximating the indicator function by a delta function yields 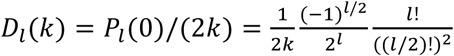. The more precise formula is given in the appendix, but the major aspects of the formula are easily seen from the last expression: the sampling thins as the reciprocal of the Fourier radius, and the absolute value of the kernel stays bounded away from 0, for all *l* even. Two graphs related to *D*_*l*_(*k*) are shown in Supplemental Figure 1 and Supplemental Figure 2A.

### 3.4. Thinning of the Sampling Function and Investigating Zeros

In supplemental Figure 2, we plot the product *k* D_1_(k) as a function of *k*, for the kernel given by (B.10). For the delta function approximant, given above, the product is just P_1_(0)/2 which is a constant independent of *k*. The full expression for *k* D_1_(k) asymptotes to this quickly and for all l (although only l=4 is shown), meaning that the kernel, which maps projections to sampling, is seen to depend almost exactly on 1/Fourier radius. This implies that the patterns should be minimally affected, with the only nontrivial aspect being the possible appearance of zeros occupying Fourier space as the Fourier radius is increased. This idea is reinforced by Supplemental Figure 2B where the sampling plots are generated at Fourier radii (FR) 20, 60, and 180. For example, at thrice the distance from the origin, there are 9 times as many sampling points on the sphere, but the number of points in a plane distant from the origin is only thrice the original. Therefore, we expect that the sampling decreases as a factor of 3. Indeed, the patterning of the sampling for FR=60 is identical to the patterning for FR=20, but with only one third the sampling across the surface of the sphere. Likewise, a histogram of the sampling values over the points on a sphere shows a similar pattern, but with only one third the sampling (value along the X-axis). Analogous behavior is observed for the case with FR=180. For “top-like” views, a similar behavior is observed (Supplemental Figure 2C). The one behavior that we observed is, when there is an excessive number of zeros in the transform, the SCF* values tend to increase slightly with increasing Fourier radii. The detailed behavior of SCF as a function of Fourier radii may depend slightly on lattice type and other non-universal features, and is therefore beyond the scope of this work. The user is able to re-enter the Fourier radius at which the sampling is determined via entry box on the GUI. in this way, the patterning of the sampling can be compared as the Fourier radius is increased. As a default, the user can begin with a Fourier radius that is ½ the boxsize of the dataset, then adjust down. A small Fourier radius should provide a similar sampling pattern, but will be more computationally efficient to process. To summarize: 1) the patterning stays approximately the same regardless of the Fourier radius, although the number of zeros will change, 2) the functional behavior depends only on the inverse Fourier radius, and 3) the information on a single spherical shell of Fourier space is equivalent to the information in the patterning of the original set of projection directions. Mathematically, these are nearly equivalent statements.

### 3.5. Sampling in a Fibonacci spiral on the sphere

For the purposes of plotting the sampling restricted to a sphere of Fourier radius, *K*, a Fibonnaci spiral is created with a number of independent points that should be attributed to such a hemisphere: namely, 2*πK*^2^. If the uncertainty of angular assignment is *δθ*, this should correspond to an approximate maximum angular frequency of: *L*_*δθ*_ = 1/(*δθ*). Now the maximum (even) angular frequency, *L*_max_, that are supported on a sphere with 2*πK*^2^ points, can be found by equating this last number by the number of points that are described by all even frequencies up to *L*_max_, which is (*L*_max_ + 1)(*L*_max_ + 2)/2. Ideally, we would like

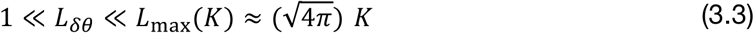

The last equality is the approximate solution for *L* for the previous quadratic when *K* large. Simply put, we should ideally like to sample the sphere more finely than the uncertainty in the angles (or rather more finely than the fluctuation in the patterning of the angles on the projection sphere). Conversely, we would like to smooth the projection directions (which lowers *L*_*δθ*_) relative to the sampling on the sphere. When evaluating the sampling on the spiral, each point is checked against the sampling condition (3.1) and the total number of occurrences of the sampling condition is tallied. The resulting integers are recorded, resulting in the vector of integer sampling along the spiral.

For a good value of analysis for the Fourier radius, a final inequality emerges from the consideration that we would like the amount of sampling at that radius to still be large on average. Therefore 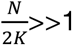. Combining this with (3.3), we should like:

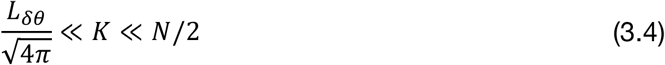

Although we have not studied the issue extensively, for typical reconstructions *K* taking values between 10 and √*N* should show patterning of the sampling to be approximately the same. Even if the angles are determined to a high accuracy, the patterning of the density of projection views is typically rather smooth, meaning that the first term in (3.4) is usually not large. The implication is that we may not need very large Fourier radii (far below Nyquist) in order to see the correct patterning.

### 3.6. Calculating the SCF and SCF*

The SCF measures the expected decrement in the SSNR as it depends on the set of projection views, keeping all other aspects of the problem the same. For the spiral described in the last section, there is a one-dimensional array of numbers, Sp, that keeps track of the count of the sampling occurrences on the surface of the sphere. If all of the values of sampling are non-zero (which is typically the case), the average of these positive numbers may be calculated as well as the average of the values of the reciprocals. The product is taken, which is a number that must be larger than unity (see Appendix of Baldwin & Lyumkis [8]). The reciprocal of this number is the SCF. In formulae: 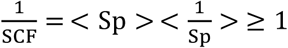, if there are zeros in the sampling array, then one must estimate how the correctly formulated SSNR (which we term SSNR* and has variance assigned to unmeasured values), is decremented due to the projection views, as described in [8]. This decrement is termed SCF^*^. If the average of the non-zero values of the reciprocal sampling is augmented by the ratio of the number of zero to non-zero values, then the product with the average sampling will again be a number that can be no less than unity. This product is 1/SCF^*^, and SCF^*^ therefore is lower than one. The inverse of SCF* can be written as the average sampling of the occupied sites, times the weighted average of the inverse sampling and unity. In formulae: 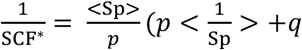. 1) ≥ 1, where *p* = 1 − *q*, and *q* is the fraction of unmeasured sample points (see [8]). The simplest example is where the *p* measured fraction all have the identical value of sampling, *v* ≥ 1. Then one calculates: 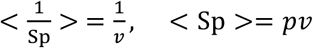, and 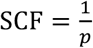, SCF^*^ = 1/(1 + (*v* − 1)*q*). Note that SCF > 1, but SCF^*^ ≤ 1. Indeed, we could have a situation where only a small part of space is sampled (*p*<< 1), but that part is sampled very highly (*v* ≫ 1) a situation that we have termed “hyper-sampling”. In that situation, SCF ≫ 1, but SCF^*^ ≪ 1. Nonetheless, the inhomogeneity in sampling drives the correctly formulated SSNR lower, as implied by SCF* < 1.

### 3.7. Sampling and Symmetry

In this sub-section, we describe how to formulate an expression for the sampling, when a symmetry is specified. If the rotation of a map acting on a model is given by = *Z*_*ψ*_*Y*_*θ*_*Z*_*ϕ*_, then the position on the projection sphere may be argued to be

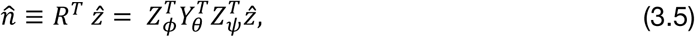

which is the application of the transpose of the map transform (see Baldwin, Lyumkis [B.13]) to the unit vector 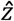. Note that the last Euler angle, *ψ*, is irrelevant to the product, which is why we can parameterize a point on the projection sphere by the first two Eulers.

Euler angles may fall within the asymmetric unit of some Platonic symmetry. Therefore, we might wish a single projection to represent itself as well as those projections associated to it by symmetry. Therefore, on the GUI there is a dropdown menu to select symmetry C, D or Platonic solid symmetries. If a representative symmetry is given by a matrix operation Sym, then the original rotation, *R*, yields equivalent copies, *R* Sym, with Sym being applied first because the symmetry pertains to the map, and not to the situation after rotation. Now, points on the projection sphere are given by:

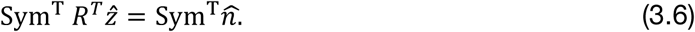

Since the full set of symmetries is given by 2*E* where E is the number of edges, we get a proliferation of unit normal by a factor of 60, 24 and 12 for icosahedral, octahedral and tetrahedral symmetries respectively. Once the symmetry is chosen, the extra unit vectors are created, and the sampling is created as described above in sub-section 3.4. Typical application of these symmetries give much more complete sampling and higher values of SCF (or SCF^*^).

### 3.8. Geometry of Tilting

In this sub-section, we describe how tilting the grid gives rise to new sampling, and a heuristic derivation of the type of patterning that is observed. The application of a tilt is to apply an additional Euler angle to the string of operations listed in Eq (3.5), and which we can assume to be with respect to the y axis: *Y*_tilt_. This will change the effective Euler angle of a given particle to a rotation given by: *Y*_tilt_ *R*, with the ultimate point on the projection sphere given by:

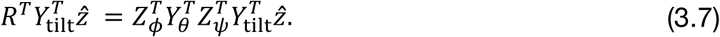

This is how the projection is plotted after choosing a tilting angle with the tilt slider within the GUI. Notice that the last Euler angle, *ψ*, which is irrelevant when plotting original projection directions, needs to be incorporated to calculate the new projection direction in (3.7). Once the new projection direction is calculated, the sampling can be determined in the same manner as before.

#### 3.8.1. Uniform in-plane rotation

To understand the patterning that results from tilting, it is an instructive exercise to show that tilting maps points on the projection sphere to circles centered around these points, under the assumption that the last in plane rotation is uniformly distributed before the action of tilting. Although, this is never strictly the case in practice, this reflects the general appearance that is observed (points mapped to circles, due to tilting). To see this mathematically, we can take the inner product of these points which will yield

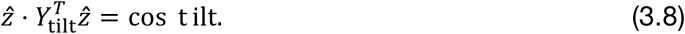

This does not depend on the last in plane rotation, which we are using as a control parameter, which we can vary. The new position on the projection sphere is at a fixed angle with respect to the original position. To understand further the meaning of the expression (3.7) when *ψ* is randomized, it is easiest to evaluate at the top of the projection sphere, when the first two Euler’s are zero. In that case:

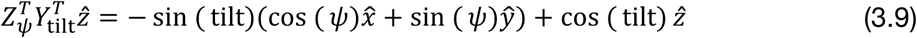

As *ψ* is swept through a full cycle, (3.9) describes a circle around the top of the sphere at the tilt angle. The situation is the same no matter what are the first two Eulers (the same sort of argument holds). The locus given by (3.9) as *ψ* sweeps from 0 to 2*π* is a circle of points at the angle “tilt” with respect to the original projection direction. The assumption that the particles have random in-plane rotation usually holds for almost all cryo-EM specimens – however for cases that this might not be true (perhaps due to biochemical tethering to a anisotropic substrate like functionalized graphene oxide), this assumption will have to be reconsidered.

#### 3.8.2. The tilting transform, under uniform last Euler, may be diagonalized via harmonics

The tilt maps a single Euler angle or point on the projection sphere to a separate unique point. However, if the last Euler is randomized, (as described above) then, if 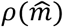 is the (possibly discrete) density of projections at the point on the unit sphere, then

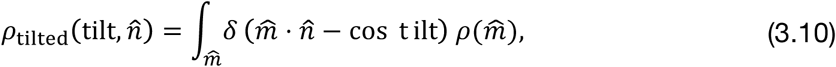

that is the tilted distribution of projections is formed from all unit normals that are at the angle given by the tilt from the original normal. When there are a great many projections, this is very nearly the case.

Similar to the arguments in Sec 3.7, this can be seen to be an angular convolution (the kernel depends on the angular distance between normal vectors). To diagonalize this transformation one can go to harmonic labels, as described before, and look at the projection of equation into the *L*^th^ harmonic subspace (the functions have an inversion symmetry so we do not need to consider harmonic subspaces of odd degree):

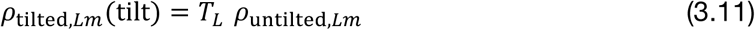

where *T*_*L*_ is the kernel implicit in Eq. 3.8: *T*_*L*_ = *P*_*L*_ (cos t ilt) (see appendix). Generally speaking, the act of tilting, in such a scenario, on the sampling can be formally undone, and might be used in inferring the original set of orientations in reconstructions performed after tilt.

## 4. Discussion

One of our major goals is to begin quantitatively evaluating angular sampling distributions and their effects on cryo-EM reconstructions. To this end, the current work introduces surface sampling plots as a complement to the conventional Euler distribution plots. To the best of our knowledge, little attempt has previously been made to directly interrogate Fourier space coverage, including unsampled regions, for pure single-particle reconstructions, although the issues ([29, 30] including point spread functions) are discussed in tomography and single particle tomography [31-33]. Surface plots of the sampling provide a quantitative measure of Fourier space coverage along particular directions and highlight regions of sampling modulation. As we will make clear in subsequent work and which is already intuitive [14, 34-36], regions that are over/under-sampled will have a direct relationship to increased/worsened resolution in the appropriate direction. Therefore, the surface plots provide an intuitive understanding of the degree of orientation anisotropy, which is not always apparent from Euler distribution plots. Lastly, zeros within Fourier space are observed in the plots and highlighted in our representations. Their presence should be a red flag to the user and would suggest that alterations to the data collection scheme, e.g. through tilting the grid, is warranted.

The GUI provides a means by which to quickly evaluate an orientation distribution, the effects of tilting the grid and estimating the resulting effect on the modified orientation distribution, Fourier coverage, and the SCF/SCF* values. The workflow for the GUI is simple and requires only a three-column angular orientation file containing the angles, Ψ, θ, and ϕ. Parameters like symmetry and Fourier radius may be altered by the user. If there are regions of Fourier space that are unfilled, one may also interrogate for how the fraction of zeros increases as Fourier radius is increased. The slider bar allows the user to obtain immediate feedback illustrating the effects of tilts. This can be beneficial during data collection in real time, as the user can immediately adjust the data collection scheme according to some optimal tilt angle. Although the simple conception of the sampling function may not necessarily provide a precise approximation to the reconstruction weights, it is useful on two accounts: 1) to model the accrual of sampling, and therefore signal; 2) to identify empty regions of Fourier space, where we could anticipate the decrement of SNR and the rapid (singular) increase of SNR, when sampling is reintroduced. We advocate it as a useful construction for visualizing the patterns of Fourier space that are filled. In summary, our tool: (1) Enables visualizing the Fourier space sampling constrained to a sphere, yielding a histogram of sampling values and plotting the fraction of empty regions of Fourier space as a functions of Fourier radius; (2) Returns the SCF metric for these sampling distributions, and SCF* for sampling distributions with empty regions; (3) Enables visualizing the possible effect of symmetry on these metrics; (4) Enables determining the effect of tilting on the sampling and on the same metrics. We envision that future improvements will build upon this baseline framework established here.

There is a remaining problem in the current application of the SCF to experimental data. Since the orientations are not known *de facto*, we currently assume that they are properly and correctly assigned during refinement. In other words, the SCF is calculated from a given distribution, regardless of whether the provided orientations are correct. This need not be the case. Orientations can be slightly or even severely mis-assigned. Furthermore, noisy cryo-EM datasets almost always have some particles interspersed throughout Euler space. Random distributions will elevate the absolute value of the SCF, and they will reduce the number of zeros in the transform. Thus, there are currently several major questions that are subject to ongoing investigation: how and to what extent do false positive orientation assignments elevate the SCF, and therefore misleadingly elevate SSNR? How often are zero values in the transform encountered in experimental cryo-EM reconstructions? Since zeros attenuate the SCF, and, in principle, should be encountered within “top-like” reconstructions, what is the effect on the SCF and SSNR of masking zeros with false positive assignments? How does tilting affect experimental cryo-EM datasets? Can an optimal till angle be determined for a given sampling distribution?

Cryo-EM reconstructions are appearing at an ever-increasing rate and are being archived within the electron microscopy databank (EMDB) [37, 38]. The EMDB summarizes important metrics relevant to the experimental reconstructions. Depositions include information about the experiment, the microscope, statistics on data collection, details of the employed software, strategies for image processing, and other important parameters. An important set of statistics pertains to common quality control metrics for experimentally-derived reconstructions, including the FSC. However, none of this information includes a description of the types of views that were acquired or can reflect any anisotropic aspect of the collection, which remains one of the most common problems in the field. As noted before, there is typically preferred orientation in cryo-EM [6] and the sets of particle views obtained can vary widely from the investigation of one protein to another, and even from one collection to another of the same protein. None of the extant metrics summarize these issues. Surface sampling plots help quantify the effects of a given orientation distribution on Fourier space coverage. The uniformity of sampling plots is determined with the SCF (or SCF*) value. We propose their use as a complementary analytic to conventional Euler plots. In the future, it would be necessary to determine their utility in experimental cryo-EM datasets and to establish a relationship between SCFs and 3D FSCs. These efforts will help the field better understand the effect of non-uniform sampling on both global and directional resolution.

## 5. Acknowledgements

PRB and DL would like to thank David DeRosier for discussions. PRB and DL are supported by grants from the NIH: DP5 OD021396, R01 AI136680, and U54GM103368.

## 6. Data Availability

The code associated with this work is available at: https://github.com/LyumkisLab/SamplingGui

## Appendix

### A. Plotting the projection directions on the plane

This refers to the plotting of the projection directions, which we introduced mathematically in section 3.1. If we make a scatter plot of points (*x*_*j*_, *y*_*j*_), we should ask how best to draw them to reflect their original packing on the surface of the sphere (*x*_*j*_, *y*_*j*_, *z*_*j*_) with each of the last vectors, *j*, normalized to unity. One can write, in spherical coordinates, the original (*x, y, z*) = (sin *θ* cos *ϕ*, sin *θ* sin *ϕ*, cos *θ*). We would like to plot (*x, y*) to have the same measure as uniform measure on the sphere, which is sin *θ* d*θ* d*ϕ*. If we set up the problem so that the azimuth is identical to the original value (that is *ϕ*), then

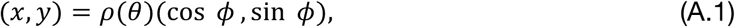

the 2D measure of these points would be *ρ*′(*θ*)*ρ*(*θ*) d*θ* d*ϕ* Therefore we equate

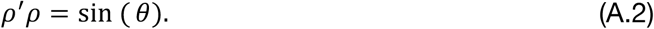

The unique solution to this ode with *ρ*(0) = 0 is *ρ*(*θ*) = constant sin (*θ*/2). This completes the discussion of section 3.1.

### B. Funk Hecke Theorem

In this appendix, we want to describe how to deconvolute angular convolutions of the sort given by (3.1), by means of transforming to harmonics. This allows us to make rigorous statements about how the sampling restricted to a sphere is equivalent in information to the original projection directions, and how the sampling thins as a function of Fourier radii (approximately as 1/*k*, which is consistent with the natural geometrical argument as the ratio of circumference to surface area). Consider transformations of the sort given by (3.1):

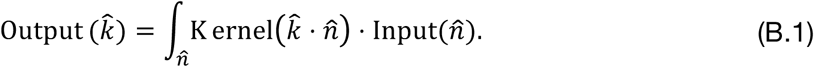

This can be diagonalized by going to harmonic labels, and looking at the projection of the equation into the *L*^th^ harmonic subspace (the functions have an inversion symmetry so we do not need to consider harmonic subspaces of odd degree).

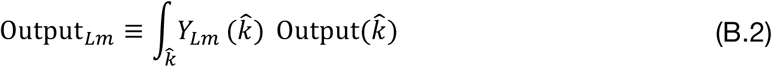

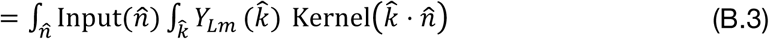

But according to the Funk Hecke theorem [28]:

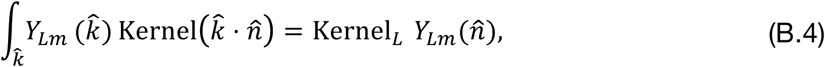

where

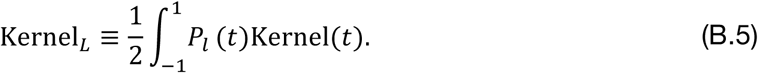

Substituting B.4 into B.3 yields

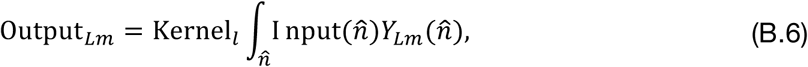

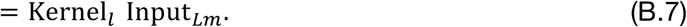

In this manner, we have diagonalized the starting equation (B.1), with an eigenvalue that only depends on the curvature label, *l*, and not the orientational label, *m*. The kernel in B.5, when applied to Eq. 3.1 becomes

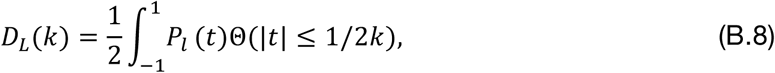

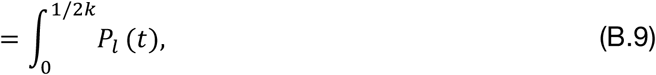

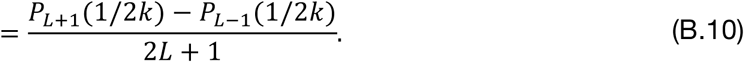

The delta function approximation to the integral is *P*_*L*_ (0)/(2*k*), and is equivalent to (B.10) in the large *k* limit. As shown in Supplementary Figs 1 and 2, the kernel, *D*_*L*_(*k*), stays bounded away from zero, and is well approximated by a functional dependence on *k*, as 1/*k*, which is the appropriate form for the thinning.

In the context of tilting, for the expressions (3.9) to (3.10), the kernel in (3.10) becomes

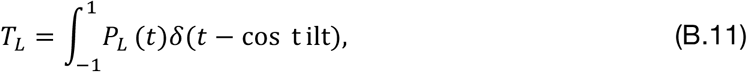

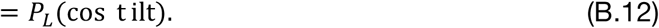

This particular expression is typically non-zero, although not strictly so: the implication being, that one can still approximately reconstruct the original sampling, from its tilted version.

### Supplemental Figures

**Supplemental Figure 1.**
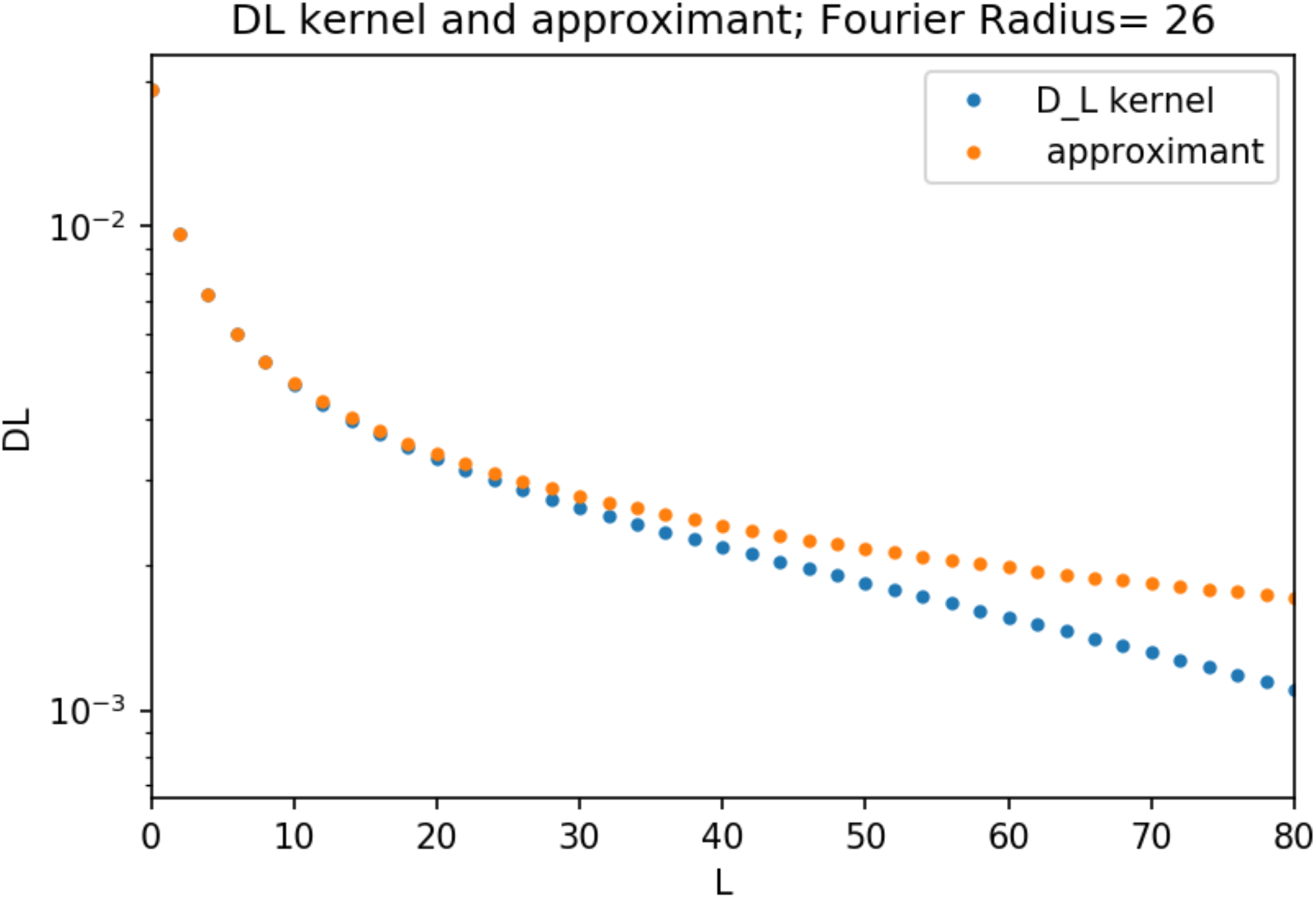
Graph of |*D*_*l*_ (*k* = 23) | (Eq B.10) and its approximant form (delta function approximation, *D*_*l*_ (*k*) ≈ *P*_*L*_ (0)/(2*k*)) as a function of curvature label, *l*. The kernel stays bounded away from zero, implying that the mapping from projections on a unit sphere to surface sampling representation is invertible.

**Supplemental Figure 2.**
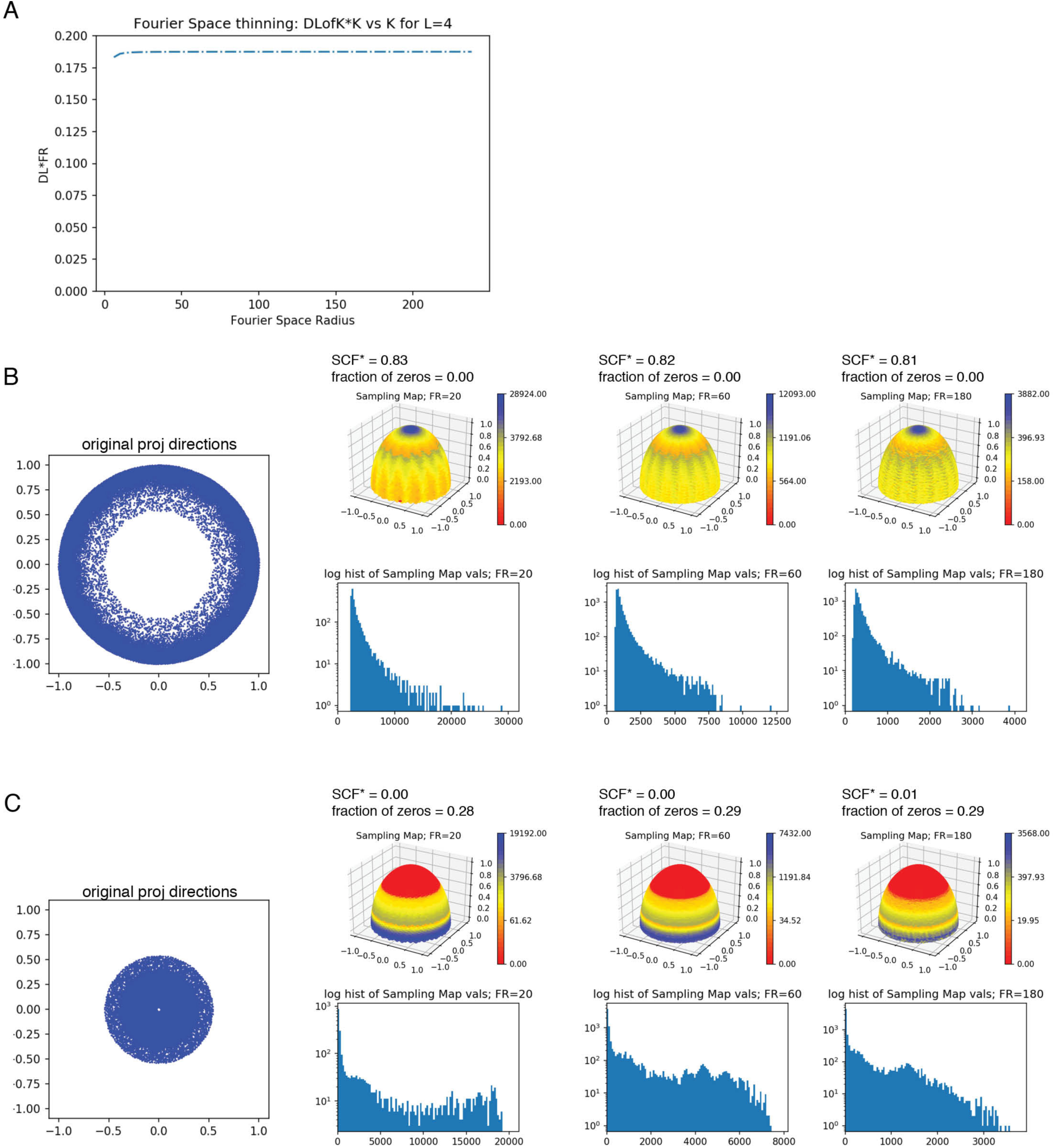
(A) Values for |*D*_*l*=4_(k)| * *k* are plotted as a function of *k*. The curve asymptotes quickly to a constant value, implying that the thinning is nearly identical to the value based on geometry, and implied by the approximant. (B-C) Thinning for experimental Euler angles of the proteasome dataset, EMPIAR-10025, separated into “side-like” and “top-like” views, as in Figure 3. Projections for the (B) “side-like” distribution and (C) “top-like” distribution are shown, alongside three sets of panels for the surface sampling plot and a histogram of the sampling values for Fourier radii 20, 60, and 180.

## Notes

https://github.com/LyumkisLab/SamplingGui

### Competing Interest Statement

The authors have declared no competing interest.

